# Ultra-low oxygen tension during *in vitro* fertilization improves embryonic and adult outcomes in mice

**DOI:** 10.64898/2026.07.10.737757

**Authors:** Cassidy N. Hemphill, Eric A. Rhon-Calderon, Alexandra J. Savage, Ana Domingo-Muelas, Christopher J. Krapp, Nicolas Plachta, Richard M. Schultz, Marisa S. Bartolomei

## Abstract

Embryo culture, a required step during *in vitro* fertilization (IVF), exposes developing embryos to altered environmental conditions not normally experienced *in vivo*, including altered oxygen (O_2_) tension. Importantly, O_2_ influences gene expression, metabolism, and the activity of enzymes that sculpt the epigenetic landscape. The lowest O_2_ tension currently used in clinics during embryo culture is 5%, despite evidence that sections of the mammalian female reproductive tract have O_2_ levels as low at 2%. Lower O_2_ may therefore better mimic the *in vivo* environment and thus lead to improved pre- and postnatal outcomes in IVF-conceived offspring. Using our validated IVF mouse model, we show embryo culture at 2% O_2_ compared to culture under 5% O_2_ significantly improves embryo cell number, the chromatin landscape in preimplantation embryos, fetal and placental development during gestation, and metabolic function in adulthood. We further uncover mechanisms by which culture under ultra-low O_2_ mediates these improvements. Overall, these results suggest embryo culture with 2% O_2_ ameliorates adverse outcomes after IVF and provide evidence that IVF could be further improved by adjusting culture conditions to model the *in vivo* environment.

## Introduction

Embryo culture, a fundamental component of assisted reproductive technologies, including *in vitro* fertilization (IVF), takes place for the entirety of preimplantation development (1). To better support embryo development, culture conditions have been continuously optimized to more accurately recapitulate the *in vivo* environment (2–4). Nevertheless, culture exposes a developing embryo to conditions not normally experienced *in vivo*, including an altered oxygen (O_2_) tension (1, 5). O_2_ plays a vital role in many cellular processes during early embryo development, including metabolic and epigenetic reprogramming (6, 7). Following fertilization, gamete-specific DNA and histone methylation must be extensively reprogrammed by O_2_-dependent enzymes such as ten-eleven translocation (TET) methylcytosine dioxygenases and jumonjiC domain lysine demethylases (KDMs) to allow eventual lineage-specific gene expression (8–12). Additionally, early embryos adapt to substrate and O_2_ availability through changes in metabolic strategies (4, 13–15). For example, the upregulation of many glycolysis related genes, including *Pfk*, *Pgam*, and *Ldha*, at the blastocyst stage enables the rapid growth and differentiation required for embryo competency (16–19). Studies in humans and several rodent species have shown the O_2_ tension is approximately 5% in the oviduct and 2% in the uterine environment (20–22).

In humans, embryo culture under 5% O_2_ tension, most commonly used by clinics today, improves blastocyst formation, pregnancy rates, and live birth rates compared to culture under atmospheric O_2_ (∼21% O_2_) (23–26). Despite these improvements, IVF with 5% O_2_ embryo culture is still associated with suboptimal fetal and placental outcomes, including pre-eclampsia, miscarriage, preterm birth, fetal and placental growth abnormalities, and a higher rate of rare imprinting disorders (27–31). Some adverse health outcomes may persist into adulthood for IVF children such as predisposition to metabolic syndromes (32–36). These findings suggest 5% O_2_ during embryo culture is physiologically suboptimal and that ultra-low O_2_ tensions (e.g. 2%) may be necessary for proper embryo development. Indeed, several limited studies in human and mouse have reported improvements to embryo development with extended culture under 2% compared to 5% O_2_ tension (37–40). In contrast, others report no significant differences in embryo development (41). Thus, further investigation is needed to clarify how ultra-low O_2_ impacts development after IVF. Additionally, molecular mechanisms underpinning effects of ultra-low O_2_ on short- and long-term offspring outcomes remain uncharacterized.

To address these gaps in knowledge, we investigated the molecular and physiological effects of 2% O_2_ embryo culture during IVF on offspring development. We utilized our validated IVF mouse model, which recapitulates key phenotypes observed in human IVF offspring (42–52). Importantly, our study spans developmental time, with offspring outcomes characterized during preimplantation, gestation, and adulthood. Our findings reveal embryo culture with 2% O_2_ during IVF improves offspring outcomes at all developmental timepoints analyzed, compared to embryo culture with 5% O_2_. These results underscore the need for continued optimization of IVF procedures to ensure offspring health.

## Results

### Ultra-low *O_2_* during embryo culture improves blastocyst morphology

We first determined the impact of ultra-low O_2_ on blastocyst formation and morphology after IVF by generating three groups of embryonic day (E) 4.0 embryos: flushed naturally conceived embryos (Naturals), and IVF embryos cultured under 2% O_2_ (IVF 2%) or 5% O_2_ (IVF 5%) (Figure 1A). To ensure consistent exposure to low or ultra-low O_2_ during embryo culture, we measured the O_2_ dynamics of culture media using a fiber-optic phosphorometer. Following a 6-h equilibration, the culture medium reached the expected O_2_ tension and maintained this tension for the duration of culture (Supplemental Figure 1A). We did not observe a significant difference in the rate of blastocyst formation between IVF 2% and IVF 5% embryos (Supplemental Figure 1B).

**Figure 1:**
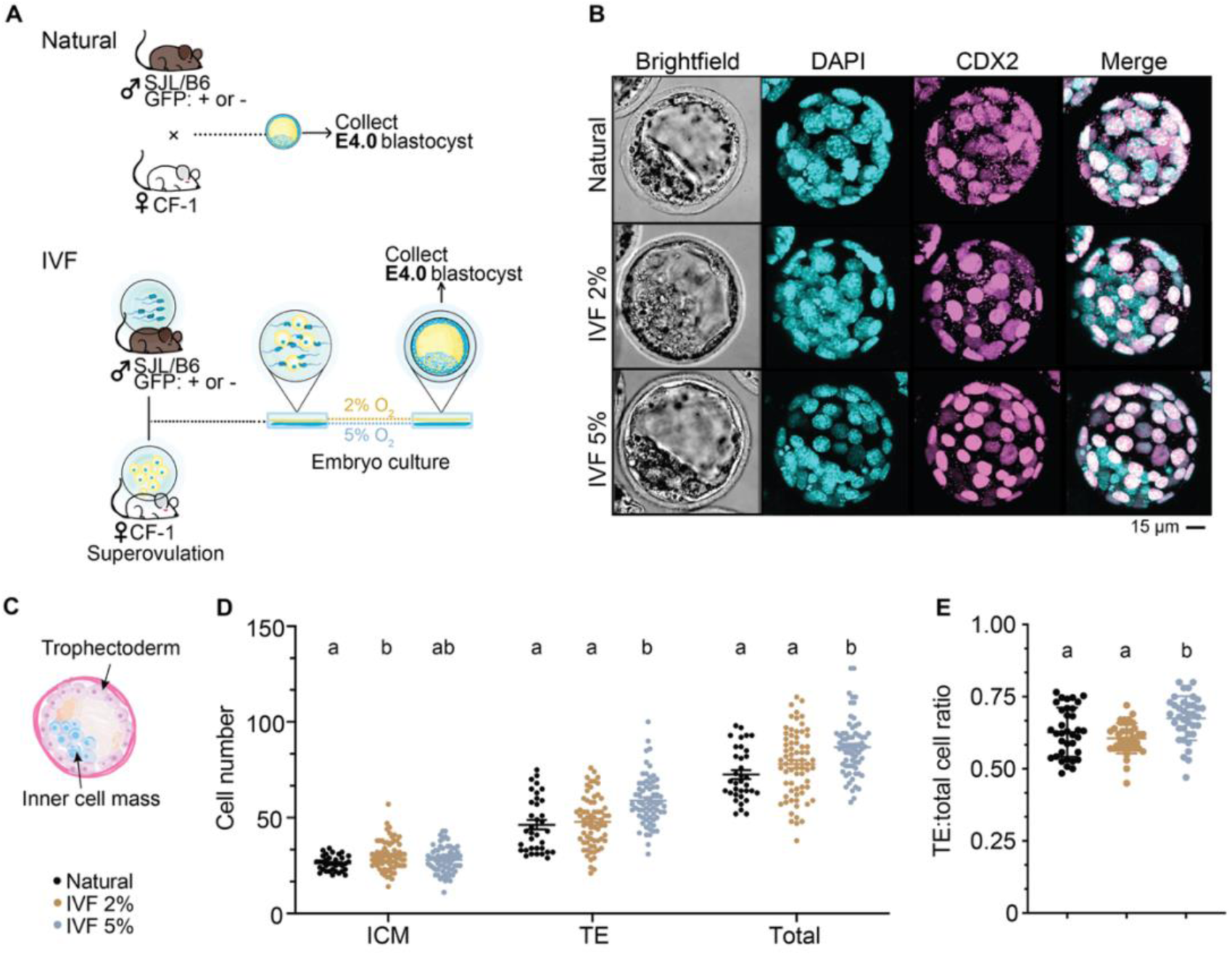
Embryo culture at 2% O_2_ improves blastocyst cell numbers and ratios at E4.0. **A**, Scheme depicting generation of Natural and IVF E4.0 blastocysts. **B**, Representative confocal images of Natural, IVF 2%, and IVF 5% blastocysts stained with DAPI and anti-CDX2 antibody. Scale bar: 15 µm. **C**, Schematic of an E4.0 blastocyst. **D**, Inner cell mass (ICM), trophectoderm (TE), and total cell counts from individual blastocysts (n ≥ 37). Cells were counted using Imaris software, with CDX2-positive cells counted as TE cells and CDX2-negative cells counted as ICM cells. **E**, Ratio of TE to total cells in individual blastocysts. Data are depicted as ± SEM and significant statistical differences between groups were determined via ordinary one-way ANOVA followed by Tukey’s multiple comparisons test. Groups with different letters denote significant differences between groups (adjusted *P* < 0.05); same letters indicate no difference was detected.

To examine embryo morphology, we stained blastocysts from each group with 4′,6-diamidino-2-phenylindole (DAPI) and an anti-CDX2 antibody to distinguish cells in the trophectoderm (TE) from inner cell mass (ICM) (Figure 1B,C), lineages that will form the placenta and embryo proper, respectively. Although IVF 2% blastocysts had a modest increase in the number of ICM cells compared to Naturals, TE and total cell numbers were not statistically different. In contrast, IVF 5% blastocysts had significantly more TE cells compared to both Naturals and IVF 2% blastocysts, driving a significant increase in both total cell number (Figure 1D), and TE to total cell ratio (Figure 1E). These findings suggest that culture under ultra-low O₂ promotes blastocyst cellular composition that is similar to Natural embryos, which may affect physiology later in development.

### IVF alters gene expression in the blastocyst

We next interrogated how embryo culture at different oxygen tensions impacted gene expression in the blastocyst. We collected single, well expanded blastocysts and performed a modified Smart-seq2 protocol. Following quality control and differential gene expression analysis with DESeq2, we identified 2490 differentially expressed genes (DEGs, 1611 up & 879 down) between IVF 2% blastocysts and Naturals, 2777 DEGs (1431 up & 1346 down) between IVF 5% and Naturals, and 1096 DEGs (751 up & and 345 down) between IVF 2% and IVF 5% blastocysts (Figure 2A). Gene function analysis revealed upregulation of cholesterol synthesis and lipid metabolism related genes and downregulation of genes involved in stress, inflammation, and apoptosis signaling in IVF 2% blastocysts compared to Naturals (Figure 2B). In IVF 5% blastocysts we saw upregulation of genes involved in actin filament organization and terms including “small molecule catabolic process”, “organic acid catabolic process”, and “carboxylic acid catabolic process” compared to Natural embryos (Figure 2C,D). Further exploration of genes belonging to these terms revealed an enrichment of genes involved in the TCA cycle and oxidative phosphorylation. After culture at 5% O_2_, blastocysts displayed downregulation of processes including chromatin remodeling, ubiquitin-dependent proteasome catabolism, and mitotic cell cycle control (Figure 2C,E), which may suggest disruption of cellular homeostasis and epigenetic patterning in these embryos.

**Figure 2:**
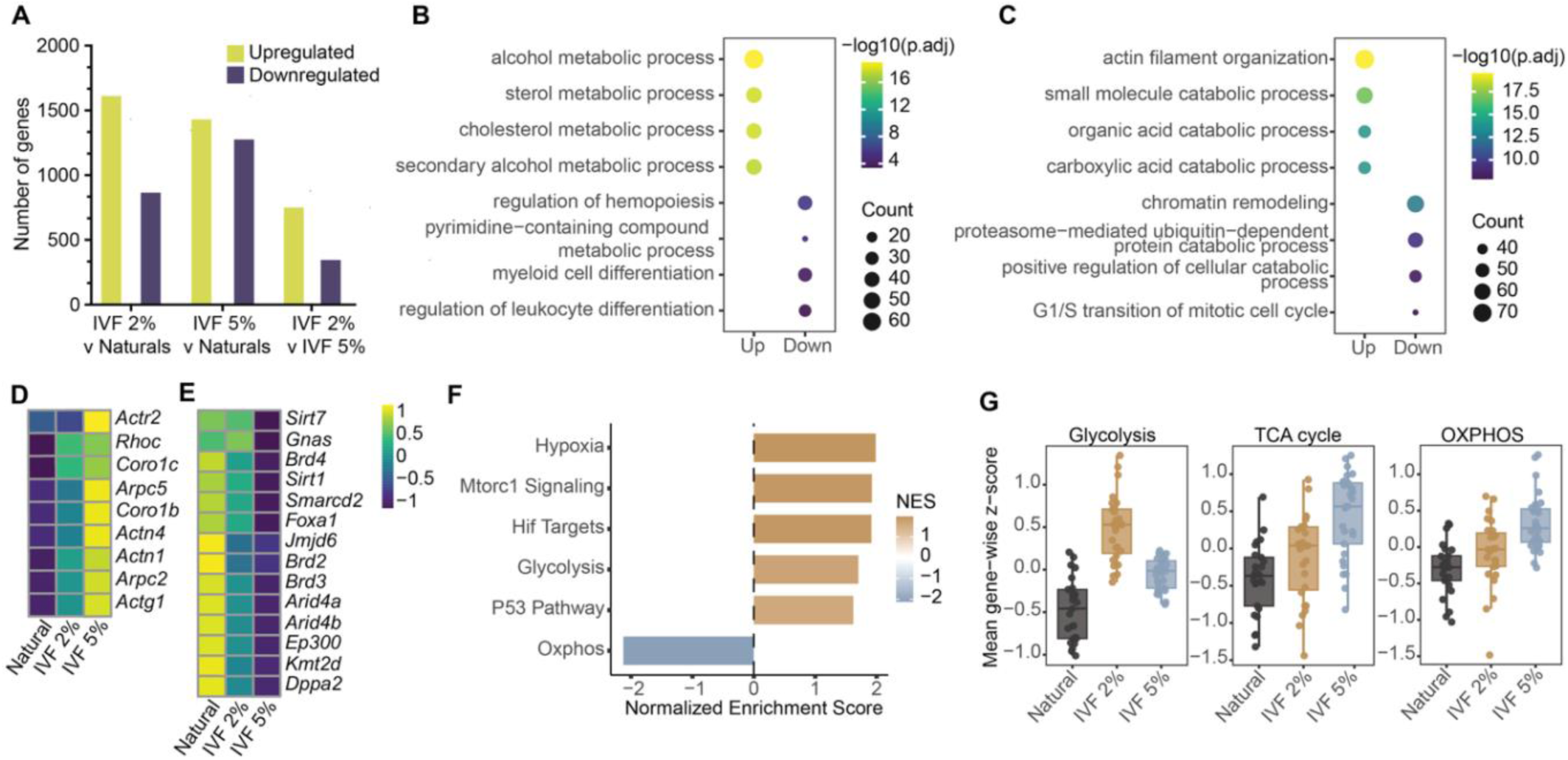
IVF alters gene expression in the blastocyst. **A**, Number of differentially expressed genes (DEGs) identified with DESeq2 between IVF 2% and Naturals, IVF 5% and Naturals, and IVF 2% and IVF 5% blastocysts. **B**, Top four enriched gene ontology pathways from up and down DEGs between IVF 2% and Natural blastocysts. **C**, Top four enriched gene ontology pathways from up and down DEGs between IVF 5% and Natural blastocysts. **D**, VST heatmap showing expression patterns of actin-related genes. **E**, Variance stabilizing transformation (VST) heatmap showing expression patterns of chromatin remodelers. **F**, Top four enriched gene ontology pathways from up and down DEGs between IVF 2% and IVF 5% blastocysts. **G**, Pathway activity scores for glycolysis, TCA cycle, and oxidative phosphorylation (OXPHOS) pathways calculated as the mean gene-wise z-score of all genes within each pathway.

Finally, a direct comparison of the two IVF groups revealed upregulation of different metabolic programs during culture, with IVF 2% embryos upregulating glycolysis-related genes and IVF 5% embryos displaying an oxidative metabolism expression signature (Figure 2F,G). Interestingly, IVF 2% blastocysts had significant enrichment of hypoxia and HIF target gene sets (Figure 2F), including genes involved in glycolysis and glucose uptake (*Slc2a1*, *Aldoa*, *Ldha*), suppression of mitochondrial oxidation (*Pdk1*, *Lonp1*, *Nampt*), ECM remodeling and invasion (*P4ha1*, *Plod2*, *Tram2*), and angiogenesis (*Egfr*, *Hey1*, *Angptl4*). These transcriptional changes suggest culture under ultra-low O_2_ activates a HIF-responsive gene expression program that supports glycolytic metabolism, tissue remodeling, and blastocyst development.

### Embryo culture at 5% O_2_ alters the epigenetic landscape in the blastocyst

Many epigenetic regulators responsible for reprogramming during preimplantation development are oxygen dependent, including DNA and histone demethylases where increased O_2_ availability enhances enzymatic activity (12, 53–55). Thus, the O_2_ environment during embryo culture likely influences the activity of these enzymes and impacts proper reprogramming. To evaluate the epigenetic status of embryos following IVF and culture, we first employed Illumina’s Mouse Infinium Methylation BeadChip to measure DNA methylation in Natural, IVF 2%, and IVF 5% blastocysts. We identified a small number of differentially methylated regions (DMRs) in IVF blastocysts compared to Naturals; 141 DMRs (1 hypomethylated & 140 hypermethylated) in IVF 2% and 77 DMRs (1 hypomethylated & 76 hypermethylated) in IVF 5% (Supplemental Figure 2A,B).

Genomic distributions of these DMRs were similar between IVF groups, with most found in intragenic regions of the genome (i.e., introns and exons) (Supplemental Figure 2C). While DNA methylation is removed from most regions of the genome during reprogramming, methylation at imprinting control regions (ICRs) is maintained. ICRs are largely involved in growth and development, and are especially sensitive to changes in methylation due to their parent-of-origin-specific mode of inheritance (56–61). Previous studies have found consistent hypomethylation of ICRs during gestation and adulthood in IVF 5% offspring tissues, including placenta, fetal liver, and adult gonads (43, 48–51, 62–64). Because the initial reprogramming period is completed in blastocysts, we tested how culture at 2 or 5% O_2_ could impact ICR methylation maintenance. To address this question, we performed targeted bisulfite sequencing for four clinically relevant ICRs; *H19/Igf2*, *Kcnq1ot1*, *Peg3*, and *Snrpn*. Surprisingly, we observed no significant changes to DNA methylation levels at these ICRs between any group (Supplemental Figure 2D), suggesting that previously reported methylation loss occurs later in development.

Histone post-translational modifications (PTMs) are also extensively remodeled following fertilization and are altered by IVF procedures (65–67). To test the impact of embryo culture at 2 or 5% O_2_ on global levels of histone PTMs in blastocysts, we performed immunofluorescence staining for H3K4me3 and H3K9me3, activating and repressive histone modifications, respectively (Figure 3A, B). Following staining and normalization to DAPI signal, IVF 2% blastocysts had a modest increase in H3K4me3 signal intensity in both TE and ICM compared to Natural embryos while IVF 5% blastocysts displayed a striking reduction in H3K4me3 stain intensity in both cell lineages (Figure 3C). We observed no differences in H3K9me3 stain intensity in either TE or ICM between IVF 2% and Natural embryos; IVF 5% TE and ICM had significantly lower stain intensity compared to the other two groups (Figure 3D).

**Figure 3:**
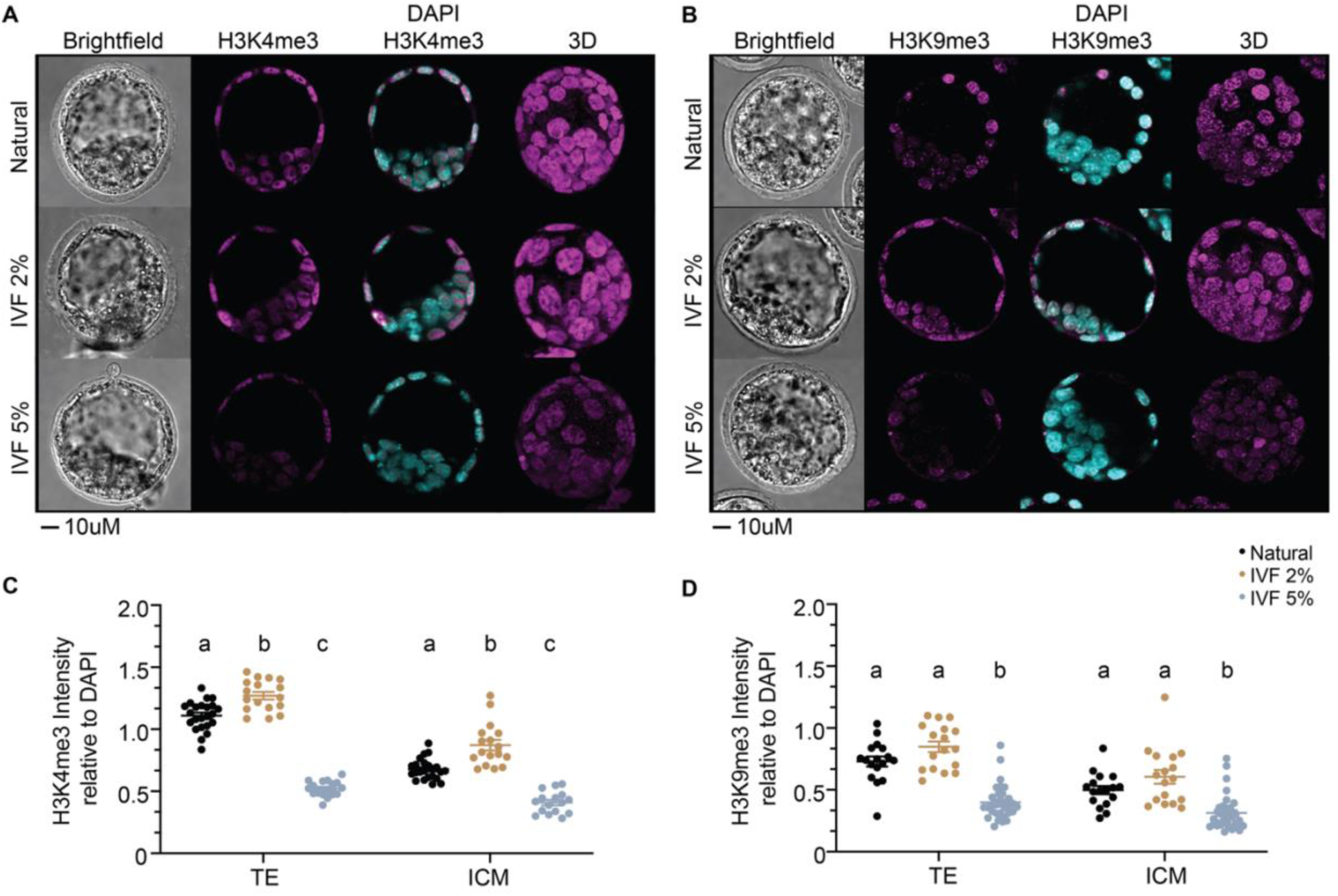
Global levels of H3K4me3 and H3K9me3 are rescued in IVF 2% blastocysts. **A**, Representative confocal images of Natural, IVF 2%, and IVF 5% blastocysts stained with DAPI and anti-H3K4me3 antibody. **B**, Representative confocal images of Natural, IVF 2%, and IVF 5% blastocysts stained with DAPI and anti-H3K9me3 antibody. Scale bars: 10 µm. **C**, H3K4me3 stain intensity in TE and ICM cells normalized to DAPI signal intensity. **D**, H3K9me3 stain intensity in TE and ICM cells normalized to DAPI signal intensity. n ≥ 16 blastocysts per group from at least two independent experiments were used for each histone modification. Data are depicted as ± SEM and significant statistical differences between groups were determined via ordinary one-way ANOVA followed by Tukey’s multiple comparisons test. Groups with different letters denote significant differences between groups (adjusted *P* < 0.05); same letters indicate no difference was detected.

Together, these results indicate that while IVF blastocysts have largely normal levels of DNA methylation, culture at 5% O_2_ drives inappropriate demethylation of both active and repressive histone PTMs. The global loss of both H3K4me3 and H3K9me3 observed in IVF 5% blastocysts, which is largely rescued in IVF 2% blastocysts, likely has consequences for subsequent epigenetic reprogramming and differentiation and long-term development.

### 2% O_2_ embryo culture improves fetal and placental outcomes at mid-gestation

Embryo culture with 5% O_2_ results in adverse developmental outcomes at mid-gestation including lower fetal weights and altered placental morphology (48, 63, 64). To determine if improvements seen in preimplantation embryos cultured at 2% O_2_ translate to better gestational outcomes, we characterized E12.5 embryos from each group. We saw a significant improvement in both fetal and placental weights in IVF 2% offspring compared to IVF 5% concepti (Figure 4A). Although we did not observe a significant improvement in fetal to placental weight ratios in IVF 2% offspring, this group had ratios that trended to be more similar to Naturals compared to IVF 5% offspring (Figure 4B).

**Figure 4:**
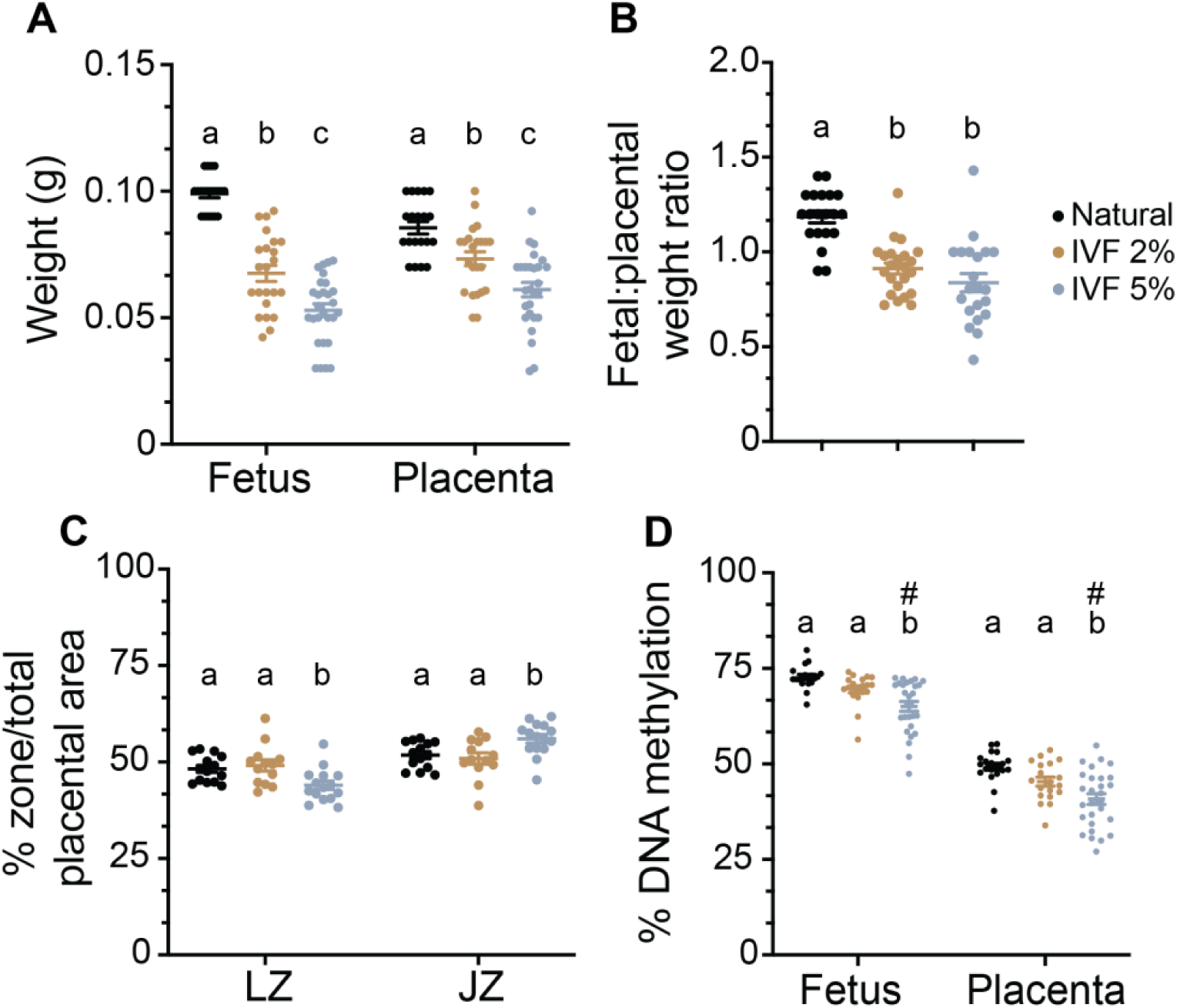
Mid-gestation offspring developmental outcomes are improved after IVF with 2% O2 embryo culture. Natural and IVF concepti were collected at E12.5. **A**, Fetal and placental weights. **B**, Fetal to placental weight ratios. **C**, Percent global DNA methylation in individual fetuses and placentas measured via luminometric methylation assay. n ≥ 20 concepti from each group were used in **A**-**C**. Data are depicted as ± SEM and significant statistical differences between groups were determined via ordinary one-way ANOVA followed by Tukey’s multiple comparisons test. Groups with different letters denote significant differences between groups (adjusted *P* < 0.05); same letters indicate no difference was detected. Variability was calculated using a Brown-Forsythe test and # represents a significant difference (*P* < 0.05) in variance.

To more thoroughly assess placental morphology, we measured placental vascularization via CD34 immunohistochemistry, a marker of fetal endothelial cells. Microvessel density was not significantly different between groups, although IVF 5% placentas trended towards a modest reduction in CD34 stain area (Supplemental Figure 3). Next, we stained a subset of placentas with hematoxylin and eosin (H&E) and measured labyrinth and junctional zone areas, regions of the placenta responsible for nutrient/gas exchange and endocrine function, respectively. We observed a significant reduction in labyrinth zone area and corresponding increase in junctional zone area in IVF 5% placentas compared to Naturals. These differences were normalized in IVF 2% placentas (Figure 4C). Overall, these data suggest that 2% O_2_ embryo culture promotes more normal placental development, potentially improving placental efficiency by preserving labyrinth zone architecture important for maternal-fetal exchange.

Embryo culture with 5% O_2_ also results in both diminished levels and increased variability in global and ICR-specific methylation in fetal and placental tissues compared to natural offspring (43, 44, 49–51, 62, 63). To determine the impact of ultra-low O_2_ embryo culture on DNA methylation at mid-gestation, we first measured global DNA methylation in whole fetuses and placentas. Notably, global DNA methylation levels in IVF 2% fetuses and placentas were improved and not statistically different from Naturals (Figure 4D). We next measured DNA methylation at several ICRs described above. DNA methylation was restored to Natural offspring levels in IVF 2% fetuses at *Kcnq1ot1* (Supplemental Figure 2E). While we did not observe significant improvement in the average methylation levels at the other ICRs in IVF 2% offspring, we did see a reduction in variability of DNA methylation at these ICRs (Supplemental Figure 2E,F). Together, these results suggest that while culture at 2% O_2_ supports proper global DNA methylation establishment following implantation, ICRs remain hypomethylated, suggesting vulnerability to other aspects of IVF procedures.

### Ultra-low O_2_ IVF improves developmental outcomes at term

To investigate the impact of ultra-low oxygen embryo culture on late gestational outcomes, embryos from each group were collected at E18.5. Importantly, we saw no difference in fetal weights between IVF 2% and Natural fetuses of both sexes (Figure 5A). Placental weights of concepti from both IVF groups and sexes were larger than Naturals, although IVF 2% placentas were significantly smaller than IVF 5% placentas (Figure 5B). These changes resulted in a reduced fetal to placental weight ratio for males and females from both IVF groups, with IVF 2% females displaying significantly improved fetal to placental weight ratios compared to IVF 5% females (Figure 5C). These findings suggest that despite persistent overgrowth, ultra-low O_2_ embryo culture promotes development of more functional placentas capable of supporting fetal growth.

**Figure 5:**
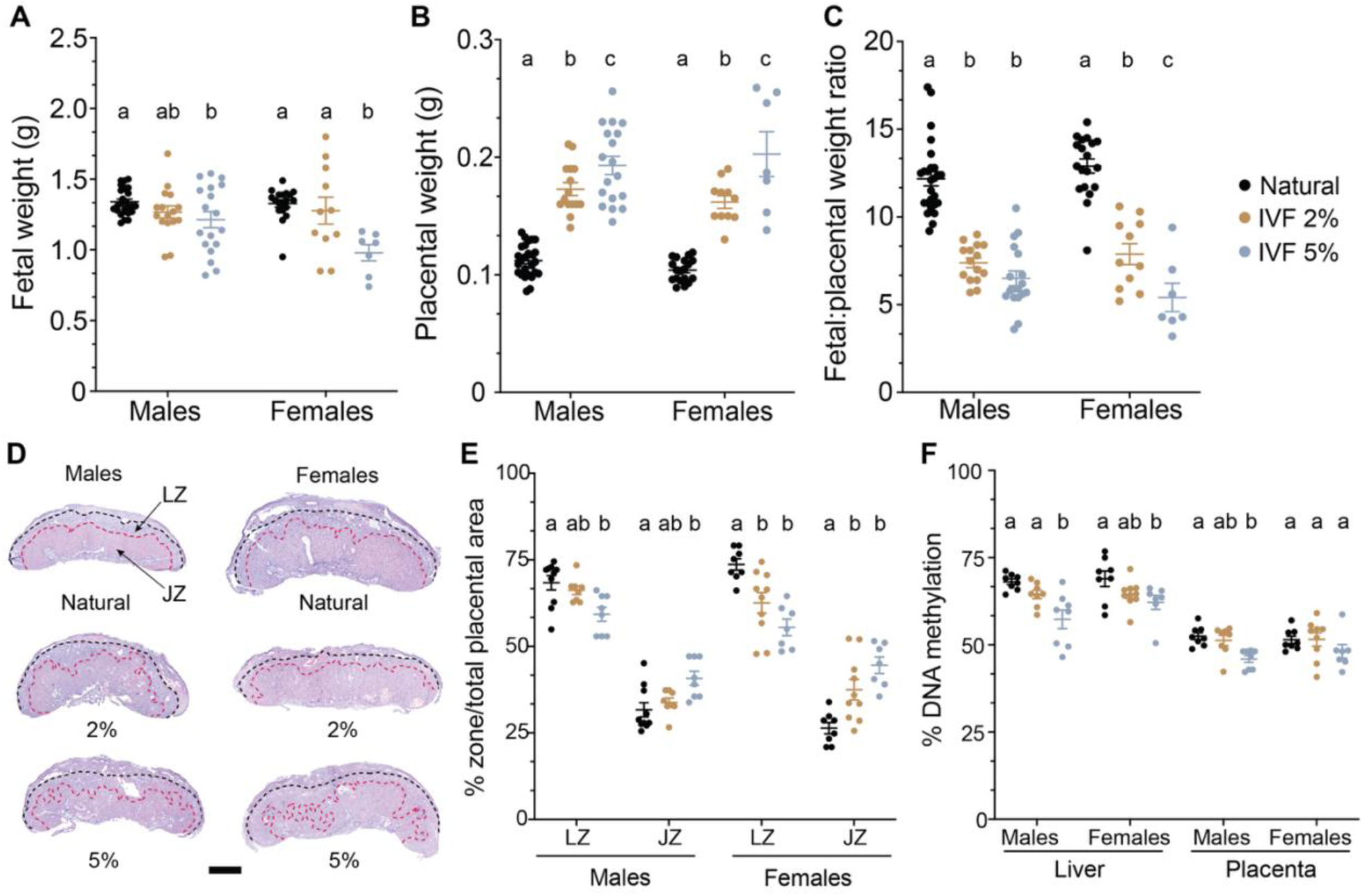
2% O_2_ embryo culture improves fetal and placental outcomes at term. **A**, Fetal weights. **B**, Placental weights. **C**, Fetal to placental weight ratios. n ≥ 16 male and n ≥ 7 female concepti from each group were utilized in **A-C**. **D**, Representative placental cross-sections stained with hematoxylin-eosin for each group and sex. The junctional zone (JZ) and labyrinth zone (LZ) are indicated by arrows on the Natural male placental section. Scale bar: 850 µm. **E**, Percentage of LZ and JZ, determined by dividing the area of each placental zone by the total placental area. **F**, Percent global DNA methylation in individual placentas and livers measured via luminometric methylation assay. n ≥ 8 male and n ≥ 7 female concepti from each group were used in (**E**) and (**F**). Data are depicted as ± SEM and significant statistical differences between groups were determined via ordinary one-way ANOVA followed by Tukey’s multiple comparisons test. Groups with different letters denote significant differences between groups (adjusted *P* < 0.05); same letters indicate no difference was detected.

To gain a more comprehensive view of placental function at late gestation, we characterized placental morphology. We stained placental sections with H&E (Figure 5E) and measured labyrinth and junctional zone areas. Interestingly, while IVF 5% male and female placentas had a significant reduction in labyrinth zone area (and associated increase in junctional zone area) compared to Naturals, we observed no difference in the areas of either zone in IVF 2% male placentas compared to Naturals. Although female placenta zone areas were not rescued relative to Naturals, IVF 2% trended to be more similar to Naturals compared to IVF 5% female placentas (Figure 5F).

Finally, we measured DNA methylation in the livers and placentas of these late gestation offspring. Globally, methylation levels in the livers and placentas of IVF 2% offspring were similar to Naturals (Figure 5G), suggesting improved global DNA methylation patterning in IVF 2% offspring tissues. In contrast, we saw largely similar DNA methylation patterns in the two IVF groups at each ICR assayed, although female IVF 2% placentas exhibited improved methylation at *H19/Igf2* compared to IVF 5% females (Supplemental Figure 2G,H). Thus, while some ICR methylation defects resolve by late gestation in IVF offspring, other ICRs remain susceptible to DNA methylation loss following IVF and embryo culture.

### Offspring derived from embryos cultured at 2% O_2_ during IVF display improved adult metabolic outcomes

IVF offspring display altered metabolism in adulthood (45, 46, 49, 63, 68–73). To address metabolic outcomes, cohorts of mice from each group were c-section delivered near term and fostered with natural littermates. These mice were weighed weekly until 12-weeks-of-age, at which point they were euthanized for additional metabolic assays. Both male and female IVF 2% offspring displayed improved body weights compared to IVF 5% offspring throughout adulthood (Figure 6A, B). Measurement of organ weights and normalization to body weights revealed a significant decrease in brain weight relative to body weight in both male and female IVF 5% offspring which was not observed in IVF 2% offspring of either sex (Supplemental Figure 3A,B). Of note, while IVF 5% females had significantly lower ovary to body weight ratios compared to Naturals, this phenotype was rescued in IVF 2% females (Supplemental Figure 3B), which may suggest an improvement in adverse reproductive outcomes seen previously in IVF 5% female offspring (51). Additionally, at 12-weeks-of-age, IVF 2% males had similar total cholesterol and triglyceride levels as Naturals, while these lipids were elevated in IVF 5% males (Figure 6E, F). Taken together, 2% O_2_ during embryo culture improves some metabolic outcomes previously observed in IVF offspring after 5% O_2_ culture.

**Figure 6:**
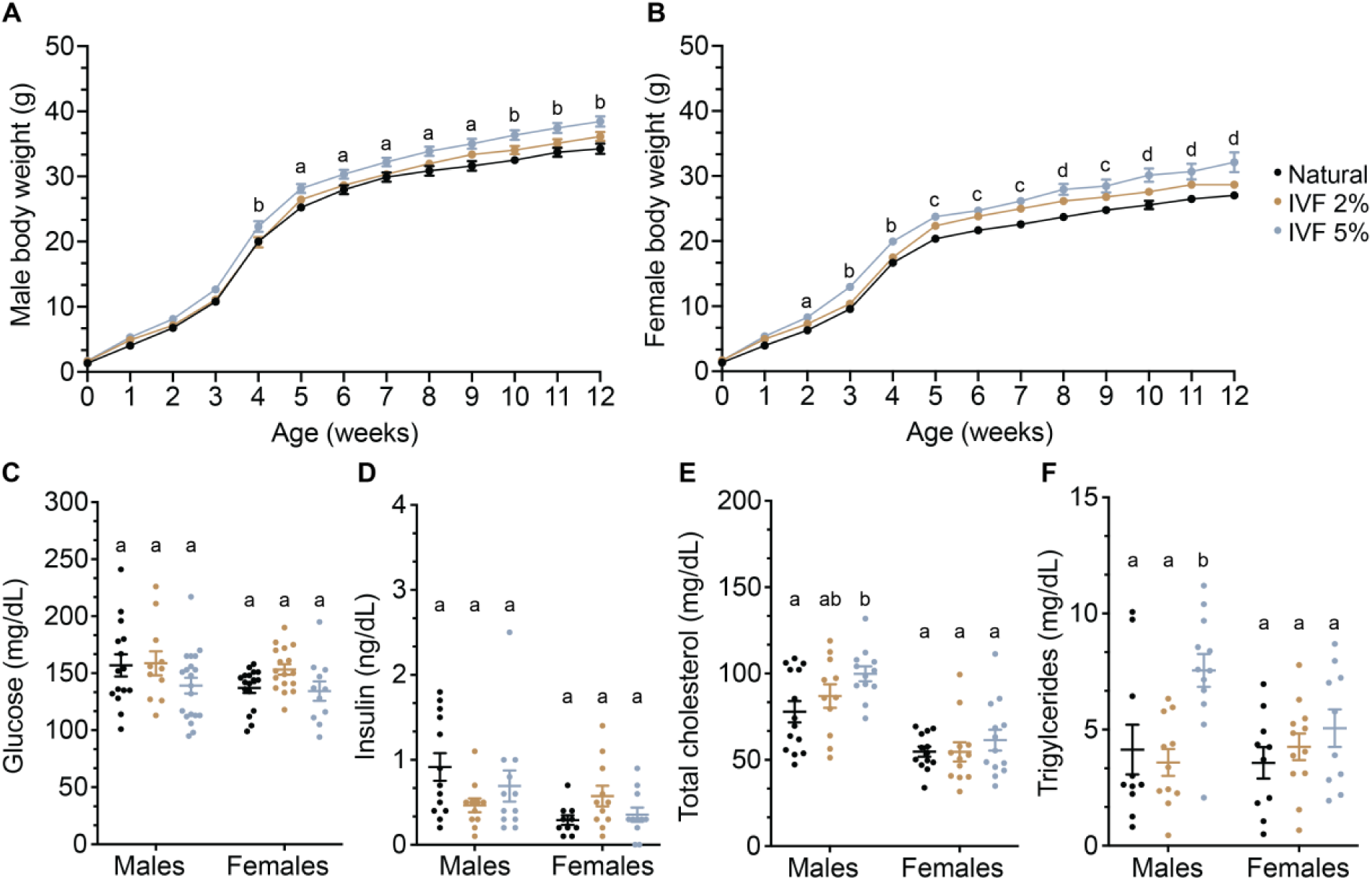
Ultra-low O2 embryo culture during IVF ameliorates metabolic outcomes in adulthood. Body weights for **A**, male and **B**, female Natural, IVF 2%, and IVF 5% offspring from birth to 12-weeks-of-age. a= significant difference between IVF 5% and Natural, b= significant difference between IVF 5% and both IVF 2% and Naturals, c= significant difference between both IVF 2% and IVF 5% and Naturals, d= significance difference between all groups. **C**, Fasting glucose, **D**, Insulin, **E**, Total cholesterol, and **F**, Triglyceride measurements using serum from 12-week offspring. n ≥ 11 male and female offspring were used in **A-F**. Data are depicted as ± SEM and significant statistical differences between groups were determined via ordinary one-way ANOVA followed by Tukey’s multiple comparisons test. Groups with different letters denote significant differences between groups (adjusted *P* < 0.05); same letters indicate no difference was detected.

## Discussion

In this study, we show that ultra-low O_2_ during embryo culture improves developmental outcomes in a mouse model of IVF. Remarkably, we observe improvements in blastocyst morphology, gene expression, and chromatin epigenetic landscape after culture at 2% O_2_ (Figure 7A), which translated to significantly improved long-term offspring health. These findings highlight O_2_’s pivotal role in shaping embryonic development and support the need for further optimization of embryo culture conditions, especially as IVF success begins to plateau (74, 75).

**Figure 7:**
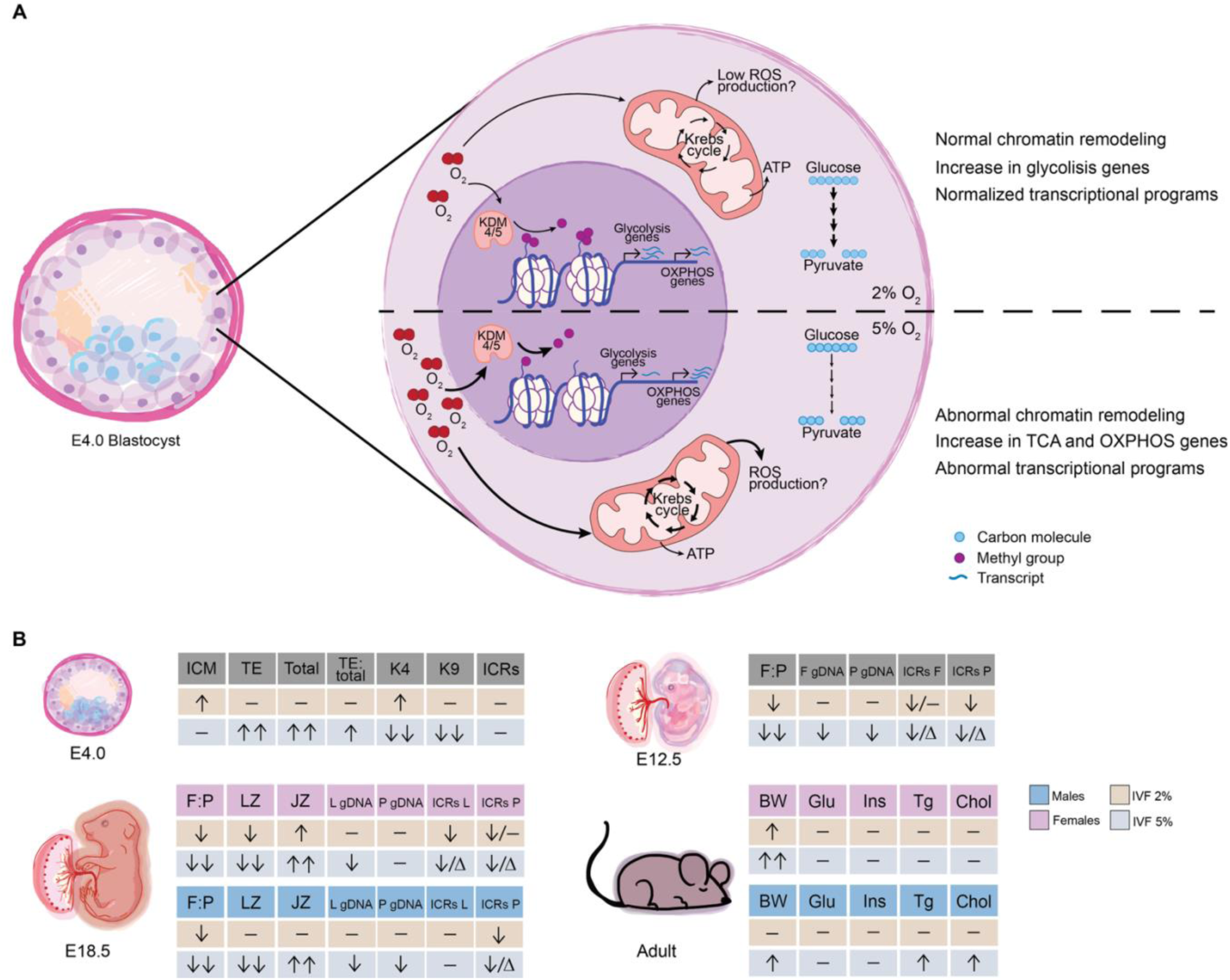
Proposed model and summary. **A**, Molecular consequences of embryo culture under 2 or 5% O_2_ in the preimplantation embryo. **B**, Summary of outcomes in IVF 2% and IVF 5% groups compared to Naturals at E4.0, E12.5, E18.5, and 12-weeks-of-age. Up arrow denotes increase, down arrow denotes decrease, dash denotes no change, delta represents increased variability.

A preimplantation embryo must undergo extensive epigenetic reprogramming following fertilization. Defects in this process are linked to congenital disorders including the imprinting disorders Beckwith-Wiedemann Syndrome and Angelman Syndrome (76–78). Moreover, mutations in epigenetic machinery, especially histone methyltransferases and histone demethylases, cause embryonic lethality in mice, as well as numerous developmental disorders in humans (79). As such, O_2_’s direct role in regulating epigenetic reprogramming has begun to be appreciated. A subclass of molecular dioxygenases, including Jumanji-C (JmjC) domain containing KDMs, utilize O_2_ as a substrate, allowing them to respond to differing levels of O_2_ (12, 53–55). We show here that embryo culture under ultra-low O_2_ preserves global levels of both H3K4me3 and H3K9me3 in the blastocyst. In contrast, culture with conventional 5% O_2_ is associated with a marked reduction in both of these histone modifications, which is in agreement with work from other groups who have observed decreased H3K4me3 in both mouse and human blastocysts following IVF (65–67). We propose that 5% O_2_, normally considered hypoxic, could actually be hyperoxic in the context of preimplantation development, with sustained availability of O_2_ leading to hyperactivity of JmjC KDMs and decreased histone methylation. Indeed, biochemical assays show increased KDM catalytic activity with higher O_2_ concentrations and reduced histone methylation; for example, the activity of KDM5A, which targets H3K4me2/3, is approximately 2-times higher at 5% O_2_ compared to 2% O_2_ (53, 55, 80). Importantly, the reduction of histone methylation observed in these studies occurred in the absence of HIF, suggesting direct oxygen-sensing by KDMs (53, 55). Further, the effect of ultra-low O_2_ on epigenetic machinery seems to be independent of changes in mRNA levels; it therefore seems likely embryo culture with 2% O_2_ promotes the normal *in vivo* activity of KDMs, leading to successful reprogramming.

Our results suggest embryo culture under 2% O_2_ supports proper metabolic programming during preimplantation development. As development progresses, an embryo must balance an increasing energy demand with the production of harmful metabolic byproducts, including reactive oxygen species (ROS). *In vivo*, this balance is regulated by substrate and, likely, O_2_ availability. While early embryos rely on pyruvate and lactate and are less metabolically active, post-compaction embryos are heavily reliant on glucose (81–84). Studies of *in vivo* developed mouse blastocysts show an upregulation of glycolytic metabolism at both the transcript and protein level (16, 19). This metabolic shift is also observed in human embryos, with proteomic analysis revealing enrichment of glycolytic enzymes including ALDOA, ENO1, LDHA, and TPI1 at the blastocyst stage compared to earlier cleavage stages (85). Consistently, we observe distinct metabolic transcriptional profiles between IVF culture conditions. IVF 2% blastocysts display expression patterns typical of the glycolytic state normally associated with blastocyst development, whereas IVF 5% embryos upregulated pathways associated with oxidative metabolism, including oxidative phosphorylation and the tricarboxylic acid cycle. Interestingly, although glycolysis is a hallmark of normal blastocyst development, glycolytic pathway activity was even greater in IVF 2% blastocysts than in Natural embryos. This may reflect an adaptive response to the *in vitro* culture environment or subtle differences in developmental progression at the time of collection. The metabolic switch observed in IVF 2% embryos may be promoted by upregulation of HIF target genes, while culture under 5% O_2_ may promote increased reliance on mitochondrial respiration during preimplantation development. Increased oxidative metabolism may be detrimental to a developing embryo, as elevated mitochondrial activity may increase ROS production (86–88). Indeed, several groups have observed markers of oxidative stress in embryos cultured at 5% O_2_, including elevated ROS, PGF2a, and DNPH, and decreased glutathione compared to *in vivo* developed blastocysts (66, 89). Glycolysis may limit oxidative stress while also supporting the rapid proliferation, biosynthetic activity, and maintenance of developmental plasticity required by a blastocyst, particularly the pluripotent cells of the ICM (90, 91). Thus, ultra-low O_2_ during embryo culture may be required to promote a metabolic program that will support proper embryo development.

In line with this proposal, we observe improved morphology in blastocysts after culture at 2% O_2_, as these embryos have TE and ICM cell counts and ratios more similar to Natural embryos. Interestingly, IVF 5% blastocysts exhibit a striking increase in TE cell number, consistent with our RNA-seq results showing dysregulated cell cycle gene expression in these embryos, and previous studies demonstrating altered cell proportions following IVF (66, 92). As TE cells are the precursors to most of the placenta, these findings could indicate very early dysregulation of extraembryonic development in IVF 5% embryos, which could contribute to the abnormal placental growth and reduced fetal weights typically observed in this group. A growing body of evidence demonstrates that placental development is a major determinant of adult health (93), with alternations in placental growth being associated with an increased risk of obesity, cardiometabolic disease, insulin resistance, and dyslipidemia later in life (93–95). Indeed, the link between placental growth abnormalities and later adverse metabolism in IVF offspring has been demonstrated in both mice and humans (45, 46, 49, 63, 68–73). Remarkedly, we show here that exposure to ultra-low O_2_ during embryo culture has lasting impacts on offspring health, with offspring of both sexes displaying significantly improved metabolic outcomes. Importantly, these improvements may be mediated by the ameliorated placental phenotypes observed in IVF 2% offspring, namely improved growth and normalized zone proportions and global DNA methylation levels.

While we provide robust evidence that ultra-low O_2_ during embryo culture improves developmental outcomes in our mouse model of IVF, we cannot be certain that these improvements will extend to human IVF. Nevertheless, mouse preimplantation development accurately recapitulates many key developmental milestones seen in human embryogenesis, including zygotic genome activation, cleavage, compaction, and blastocyst formation. Given the extended time required to reach the blastocyst stage in human embryos, this developmental difference may actually increase the vulnerability of embryos to suboptimal culture conditions. Indeed, studies of H3K4me3 in human blastocysts observed a significant reduction in this histone mark between days 5 and 6 of culture, suggesting an association between culture time and adverse effects (62, 66). Studies that have employed an ultra-low O_2_ culture strategy during human IVF have observed largely improved blastocyst formation rates and increased blastocyst quality, although these studies are limited and did not characterize outcomes after the blastocyst stage (37–39, 96, 97). Furthermore, the molecular consequences of culture under ultra-low O_2_ are yet to be defined in human embryos. While we employed a monophasic O_2_ tension culture strategy, numerous studies support a dynamic *in vivo* O_2_ environment (20–22, 98). Thus, future studies should investigate the impact of bi-phasic O_2_ embryo culture on offspring development, with an emphasis on developmental outcomes beyond preimplantation.

In summary, our work has revealed the molecular and developmental consequences of 2% O_2_ embryo culture during IVF. Our findings suggest that ultra-low O_2_ tension during preimplantation development causes early improvements in metabolic and epigenetic programming of the blastocyst, which may in turn preserve normal placental development and adult metabolic health (Figure 7B). As IVF becomes increasingly accessible and widely used, it is imperative to continue to optimize culture conditions to ensure the health of future IVF children.

## Methods

### Sex as a biological variable

Sex was considered as a biological variable for this study due to previously identified sex-specific outcomes following IVF procedures (49, 63). To address this, male and female samples were analyzed separately when possible.

### Animals

Breeding stocks of CF-1 male and female (Charles River Laboratories), C57BL/6-Tg(-CAG-EGFP)131Osb/LeySopJ male and SJL female (The Jackson Laboratory) to generate SJL/B6 male sperm donors, and CD-1 vasectomized male mice (Charles River Laboratories) were maintained in a pathogen free facility. All animals were housed in polysulfone cages and had access to drinking water and chow (Laboratory Autoclavable Rodent Diet 5010, LabDiet) *ad libitum*.

### Generation of Natural offspring

Offspring conceived naturally (Naturals) were generated by mating CF-1 females (8-12-weeks-old) in their natural estrous cycle with mature SJL/B6 males (8-24-weeks-old) heterozygous for GFP. Detection of a vaginal plug marked embryonic day (E) 0.5. Embryos were allowed to develop until E12.5 or E18.5. E12.5 offspring were euthanized and whole embryos were snap-frozen in liquid nitrogen for further molecular analysis. Placentas were cut in half across the umbilical cord: half was snap-frozen in liquid nitrogen for molecular analysis and half was fixed in 10% phosphate-buffered formalin for use in morphological studies. A subset of E18.5 offspring were c-section delivered and fostered with CF-1 mothers as previously described (50, 51, 63). Other E18.5 offspring were euthanized, livers were isolated and snap-frozen in liquid nitrogen for molecular studies, placentas were treated the same as those from E12.5 offspring.

### Generation of IVF offspring

IVF offspring were generated as previously described (49). Briefly, CF-1 females (8-12-weeks-old) were superovulated with 5 IU equine chorionic gonadotropin (eCG) followed by 5 IU human chorionic gonadotropin 46 h after eCG injection. On the day of IVF, sperm was collected from the caudal epididymis and vas deferens of an SJL/B6 male (8-24-weeks-old) heterozygous for GFP and capacitated for at least 1 h in filtered human tubal fluid (HTF; EMD Millipore) containing 3% w/v bovine serum albumin (BSA; AlbuMax, Gibco) under heavy mineral oil (Irvine Scientific). Eggs were then collected from the ampulla of superovulated females and fertilized with capacitated SJL/B6 sperm in HTF medium. After 4 h, fertilized eggs were washed in HTF medium and EmbryoMax KSOM medium containing half amino acids (KSOM+AA, EMD Millipore) before culture to the blastocyst stage in KSOM+AA under heavy mineral oil at 37°C and either 5% CO_2_, 90% N_2_, 5% O_2_ or 5% CO_2_, 93% N_2,_ 2% O_2_. After 3.5 days of culture, blastocysts were washed in warmed Multipurpose Handling Medium Complete (Irvine Scientific) with gentamicin before transfer to pseudopregnant recipients. Pseudopregnant recipients were generated by mating CF-1 females (8-12-weeks-old) with CD-1 vasectomized males 3.5 days prior to embryo transfer. Each pseudopregnant female received 10 blastocysts via nonsurgical embryo transfer. The day of embryo transfer was defined as E3.5. Pregnant females were euthanized at E12.5, E18.5 or E19.5. E12.5 and E18.5 offspring were used for molecular and morphological analysis as described above. E19.5 offspring were c-section delivered and fostered with CF-1 mothers as previously described (49–51, 63).

### Blastocyst collection

To generate Natural blastocysts, CF-1 females (8-12-weeks-old) were superovulated as described above. Following hCG injection, females were mated with SJL/B6 males (8-24-weeks-old). 98 h post-hCG injection, females were euthanized and uterine horns were isolated. Blastocysts were flushed from the uterine horns with warmed Multipurpose Handling Medium Complete. IVF blastocysts from different oxygen culture conditions were saved after 4.0 days of culture. Well expanded Natural or IVF blastocysts were either used for morphological or molecular assays.

### Single-blastocyst RNAseq

Natural or IVF blastocysts were washed in 1x Dulbecco’s Phosphate Buffered Saline (DPBS; Gibco) with 0.1% Triton X-100 (Sigma Aldrich), abbreviated as 0.1% PBT. Single blastocysts were pipetted into 5 µL 1x buffer TCL (Qiagen) with 1% 2-mercaptoethanol (BME; BIO RAD) and snap-frozen in liquid nitrogen. All samples were stored at -80°C until use.

RNAseq libraries were prepared as previously described (51, 99, 100). Library quality control was conducted using High Sensitivity DNA ScreenTape for TapeStation (Agilent Technologies) and a Library Quantification Kit (KAPA Biosystems). A final library concentration of 1.1 nM was loaded into a NextSeq 1000 sequencing machine (Illumina). Raw reads were trimmed with trimgalore and aligned to the mm10 reference genome using STAR. A count table was generated using featureCounts. DESeq2 was used to perform normalization and differential expression analysis with an FDR cut off < 0.05 and minimum fold change of 1.5. Pathway analyses were performed using the clusterProfiler R package with a p-value cutoff of < 0.05 and a q-value cutoff of < 0.2.

### Immunostaining and Cell Counts

Blastocysts were fixed in 4% paraformaldehyde in 0.1% PBT and prepared for confocal imaging as previously described (101). After blocking, blastocysts were incubated overnight at 4°C with a CDX2 primary antibody (1:1000; Abcam). Embryos were rinsed in 0.1% PBT before incubation with an Alexa Fluor 647 secondary antibody (1:500; Invitrogen). To label all nuclei, embryos were incubated with DAPI (1:100; Sigma). Embryos were imaged using a laser scanning confocal (Leica SP8).

Following imaging, 3D visualizations of embryos were created using Imaris 9.7 software (Bitplane AG). DAPI stained nuclei positive for CDX2 were counted as trophectoderm (TE) cells and those negative for CDX2 were counted as inner cell mass (ICM) cells. Total cell counts were calculated by adding TE and ICM counts. Cell counts were performed by two blinded individuals.

### Immunostaining and histone PTM measurements

Blastocysts were prepared for staining as described above. After blocking, blastocysts were incubated overnight at 4°C with a CDX2 primary antibody (1:100; Santa Cruz Biotechnology) and either an H3K4me3 (1:200; Abcam) or H3K9me3 (1:700; Abcam) primary antibody. Embryos were rinsed in 0.1% PBT before incubation with Alexa Fluor 647 secondary antibody (1:1000; Invitrogen) and Alexa Fluor Plus 488 secondary antibody (1:500; Invitrogen). To label all nuclei, embryos were incubated with DAPI (1:100; Sigma). Embryos were imaged using a laser scanning confocal (Leica SP8). Immunofluorescence images were analyzed with ImageJ software by a blinded individual.

### Low-input genome-wide DNA methylation analysis

Genome-wide DNA methylation was measured in Natural or IVF blastocysts. Samples were prepared using a modified protocol for the NEBNext Enzymatic Methyl-seq Kit. Briefly, pools of 20 blastocysts were washed in 0.1% PBT before incubation in Acid Tyrode’s Solution (Sigma Aldrich) for 2 min. After several washes in fresh 0.1% PBT, embryos were pipetted into 5 µL NEB elution buffer. Samples were snap-frozen in liquid nitrogen and stored at -80°C for further use. Samples were brought to a volume of 28 µL with NEB elution buffer and heated at 80°C for 10 min before continuing with oxidation of 5-methylcytosines and 5-hydroxymethylcytosines. Elution products after clean-up of TET2 oxidized DNA were used as input for Klenow amplification, which was performed as previously described (102). 10 µL of converted and amplified DNA was loaded onto an Illumina Infinium Mouse Methylation-12v1-0 BeadChip and run on an Illumina iScan system using the manufacturer’s standard protocol at the Children’s Hospital of Philadelphia (CHOP) Center for Applied Genomics Genotyping Core.

Raw IDAT files were processed as previously described (11, 103, 104). Processed data was analyzed using the SeSAMe package to determine DNA methylation changes between groups (FDR of 0.05, minimum of 20% differential methylation level). CG identity was assessed using the tool KYCG (part of the SeSAMe R package). Pathway analyses were performed using clusterProfiler in R.

### Placental histology and immunohistochemistry

Fixed placentas were dehydrated through a graded series of ethanol concentrations, cleared with xylene, and embedded in paraffin. Serial histological sections of 5 μm were prepared using a microtome. Placentas were stained with hematoxylin-eosin and imaged using an EVOS FL AutoCell Imaging System. Images were analyzed using FIJI/ImageJ to measure the total placental, labyrinth zone, and junctional zone area. The ratios of the junctional zone and labyrinth zone to the total placental area were calculated and expressed as percentages.

Placental sections were immunostained to detect vascular changes using the endothelial cell–specific marker CD34 as previously described (48, 63). Placental sections were scanned and digitized using the Aperio Scanscope CS-O slide scanner (Leica) at the CHOP Pathology Core Laboratory. The labyrinth area was outlined using Aperio Imagescope software (Leica), and the Aperio Blood Vessel Analysis algorithm was applied as previously described (48, 63). Data are presented as the algorithm-measured areas of CD34-positive staining relative to the labyrinth area.

### Luminometric Methylation Assay (LUMA)

Genomic DNA was isolated using phenol/chloroform/alcohol (25:24:1; Millipore Sigma), followed by ethanol precipitation and resuspension in buffer (10 mM Tris-HCl, pH 8.0, 0.5 mM EDTA). DNA (1 μg) from E12.5 placentas and whole embryos or E18.5 offspring placentas and livers was used to measure global DNA methylation by LUMA as previously described (48).

### Targeted bisulfite sequencing

Genomic DNA (1µg) was bisulfite treated as previously described(11). 50ng of bisulfite treated DNA was used to measure DNA methylation at *H19*/*Insulin-like growth factor 2* (*H19/Igf2*), *potassium voltage-Gated Channel Subfamily Q Member 1 overlapping transcript 1* (*Kcnq1ot1*), *paternally expressed 3* (*Peg3*) and *Small nuclear ribonucleoprotein-associated protein N* (*Snrpn*) via targeted bisulfite sequencing on a MiSeq sequencer (Illumina). Raw data was analyzed as previously described (11).

### Metabolic Assessment

Natural or IVF offspring that were fostered with CF-1 mothers (litters were culled to a total size of 10-12) were used for metabolic studies. Offspring were weighed weekly from 1-week through 12-weeks of age using with a calibrated digital scale. 12-week-old offspring were euthanized after 6 hours of fasting and blood glucose levels were measured via tail snip with a handheld glucometer (*ReliOn*). Liver tissue was isolated, snap-frozen in liquid nitrogen, and stored at -80°C for further molecular studies. Whole blood was collected, centrifuged at ≥ 10,000 rpm at 4°C, and serum was collected and stored at -80°C for further metabolic assays. Total triglycerides were assayed using enzymatic colorimetric assay kits from Stanbio and insulin was assayed using enzymatic colorimetric assay kits from Crystal Chem. Total cholesterol (including free) was measured using the Abcam high-, low-, and very low-density lipoprotein (HDL and LDL/VLDL) Cholesterol Assay Kit (ab65390). HDL and LDL levels were determined using Mouse HDL-Cholesterol Assay and Mouse LDL-Cholesterol Assay kits from Crystal Chem, respectively (79990, 79980).

### Culture Medium Oxygen Measurements

Oxygen measurements were performed using the phosphorescence quenching method and Oxyphor PdG4 probe (105). The PdG4 probe was diluted 1:500 in KSOM+AA for a final concentration of 0.4 µM and was allowed to equilibrate at the desired oxygen tension for 24 hours. Dishes with 50 µL probe + media drops covered in heavy mineral oil were prepared and oxygen levels were measured using a fiber-optic phosphorometer (Oxyled, Oxygen Enterprises) at various timepoints throughout a 4 day culture period.

### Statistics

All samples were statistically analyzed by ordinary one-way ANOVA or 2way ANOVA. Probabilities of genomic regions in the array were compared using a Bernoulli distribution using R v 2023.12.0+369 (R foundation for Statistical Computing; www.R-project.org/). Differences between groups were denoted as statistically significant if *P* < 0.05. Differences in variability were calculated by F test; statistical differences between groups were denoted if *P* < 0.05. All statistical analyses were performed using GraphPad Prism version 10.

### Study Approval

All animal work was conducted with the approval of the Institutional Animal Care and Use Committee (IACUC) at the University of Pennsylvania. IACUC protocol 803545 has been previously revised and approved.

## Data availability

All values for data points in graphs are reported in the Supporting Data file. Additionally, methylation array data and sequencing data included this article are available at NCBI Gene Expression Omnibus at https://www.ncbi.nlm.nih.gov/geo/, under GEO accession number GSE335256.

## Author Contributions

CNH, EARC, and MSB designed research studies. CNH, EARC, AJS, ADM, and CJK performed experiments. CNH, EARC, AJS, and CJK acquired data. CNH wrote the manuscript with edits and feedback from all co-authors. CNH is listed as first author for her conceptualization and completion of the project.

## Acknowledgments

We acknowledge the individuals who supported this work, including all members of the Bartolomei lab, Ken Zaret for use of the microtome, Jean Kim for generating the mouse model diagram, Sergei Vinogradov and Mirna El Khatib for help with oxygen measurements, and Benjamin Glass for helpful discussions and data analysis expertise.

---

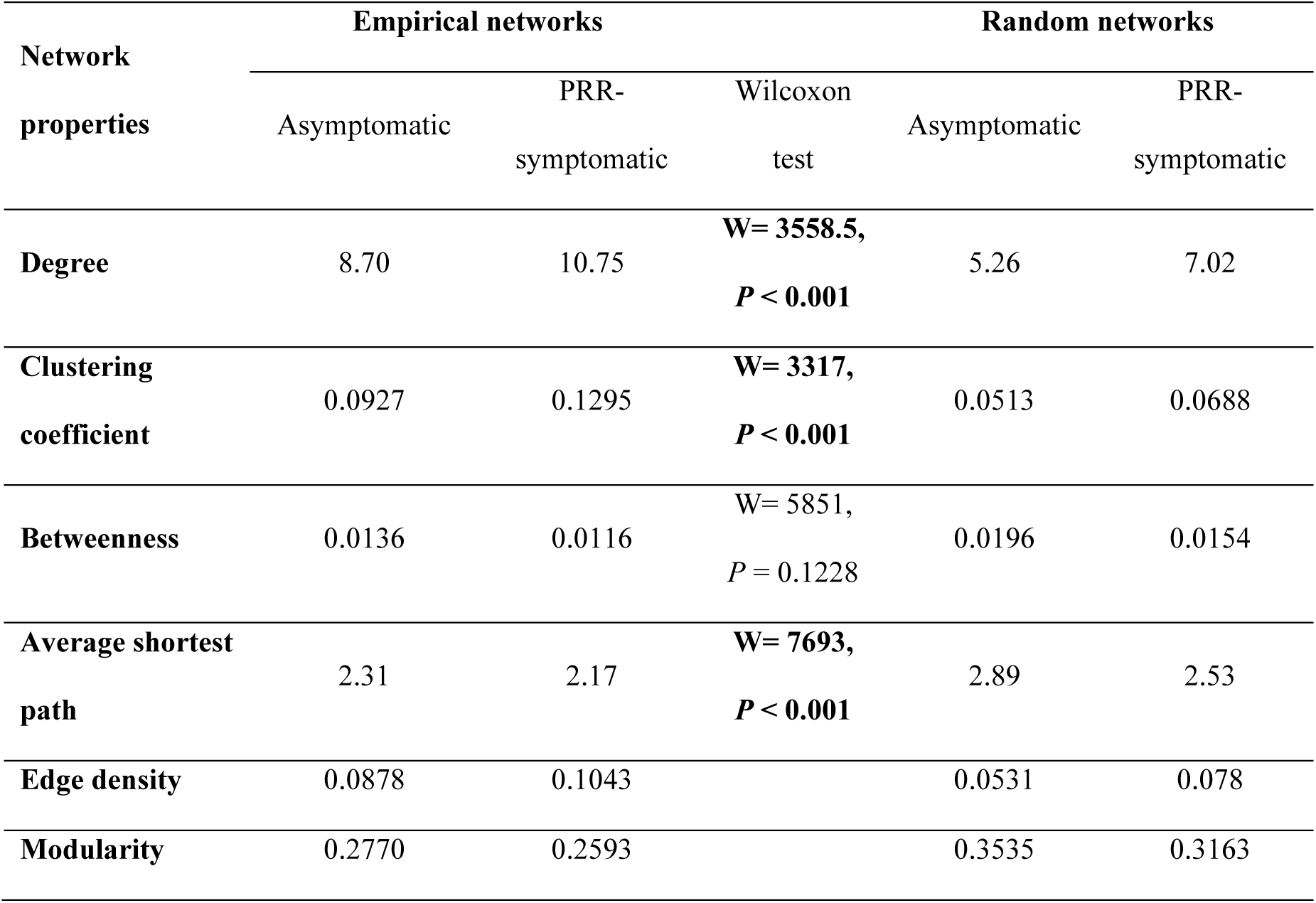

