## Supplemental figures for "Ultra-low oxygen tension during *in vitro* fertilization improves embryonic and adult outcomes in mice"

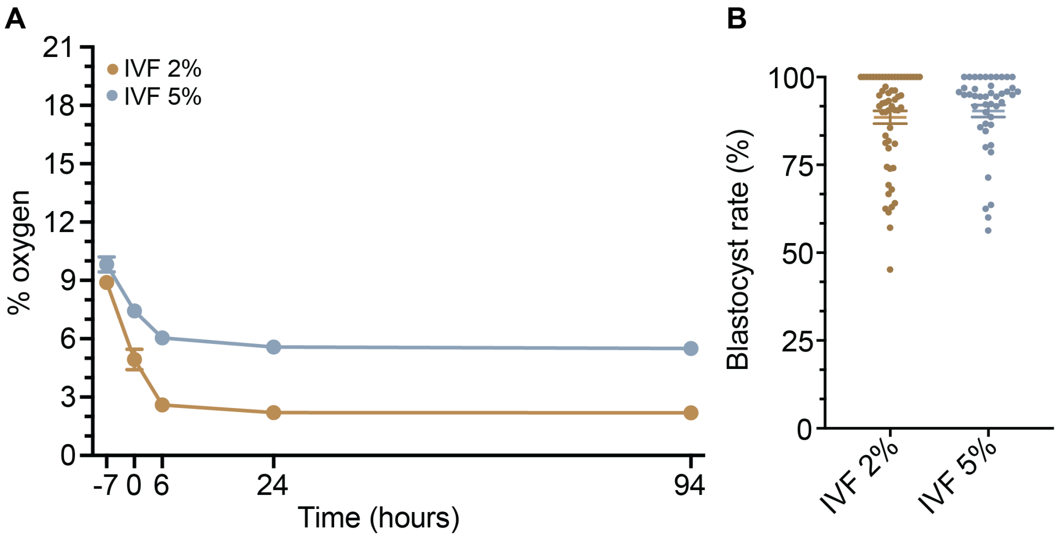


**Supplemental Figure 1: Embryo culture oxygen dynamics and blastocyst rate.**
**A**, KSOM culture medium oxygen concentration measured at -7, 0 (time at which embryos are put in KSOM media), 6, 24, and 94 hours during culture. **B**, Blastocyst rate following culture at 2 or 5% $O_2$. Data are expressed as mean ± SEM. Statistical analysis between groups was done by t-test.

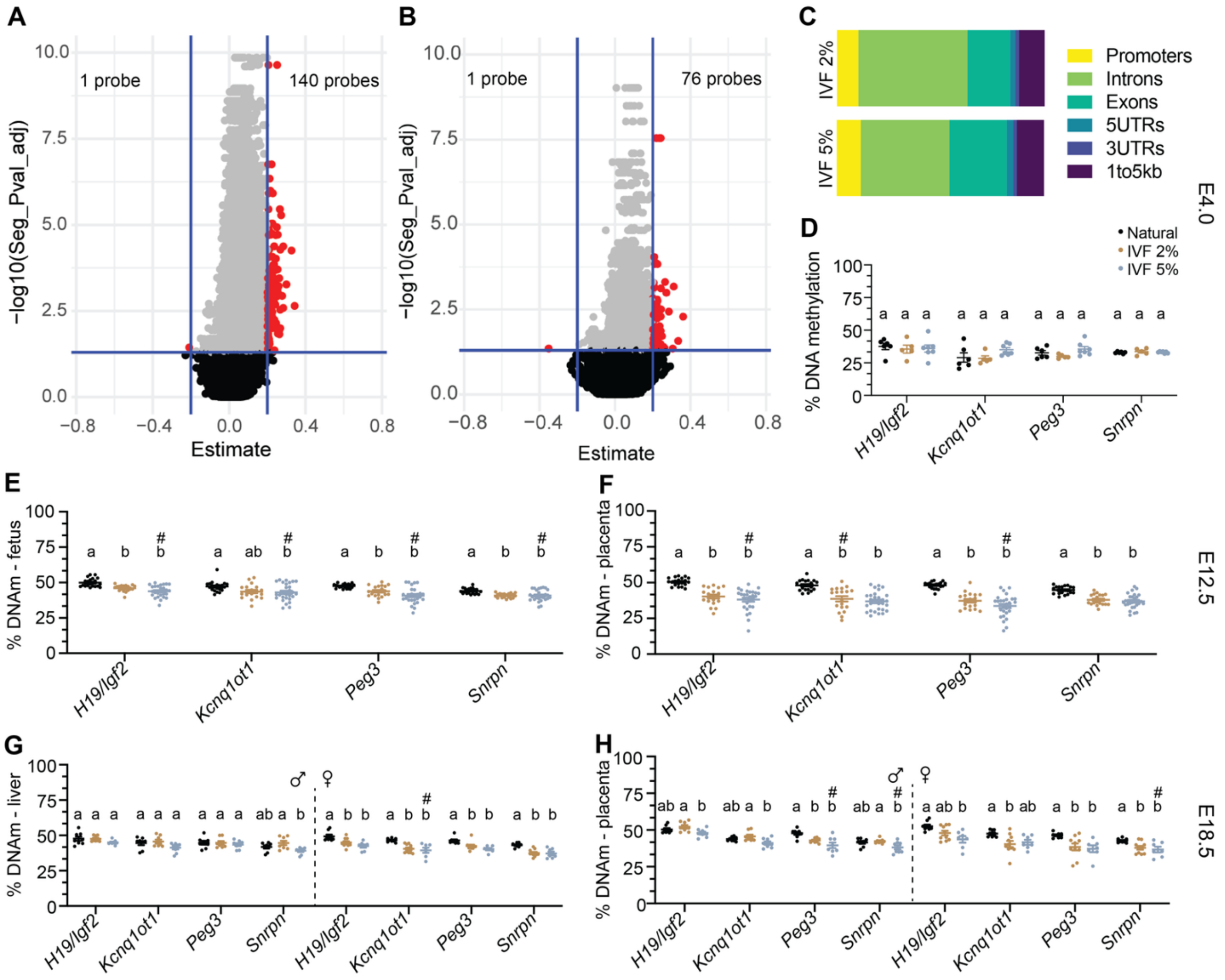


**Supplemental Figure 2: Changes in DNA methylation are seen after implantation following IVF.**

**A**, Volcano plot comparing the methylation status of IVF 2% and Natural blastocysts. **B**, Volcano plot comparing the methylation status of IVF 5% and Natural blastocysts. 20 blastocysts were pooled for each sample, and n ≥ 5 samples per group were run on the array. **C**, Genomic location of hypermethylated probes in IVF 2% (top) and IVF 5% (bottom) blastocysts compared to Naturals. Percent DNA methylation measured via targeted bisulfite sequencing at *H19/Igf2*, *Kcnq1ot1*, *Peg3* and *Snrpn* in **D**, pools of 5 blastocysts, individual E12.5 **E**, fetuses and **F**, placentas, and individual E18.5 **G**, livers and **H**, placentas. n ≥ 20 E12.5 concepti from each group were used in **E** and **F**. n ≥ 8 male and n ≥ 7 female E18.5 concepti from each group were used in **G** and **H**. Data are depicted as ± SEM and significant statistical differences between groups were determined via ordinary one-way ANOVA followed by Tukey's multiple comparisons test. Groups with different letters denote significant differences between groups (adjusted $P < 0.05$); same letters indicate no difference was detected. Variability was calculated using a Brown-Forsythe test and # represents a significant difference ($P < 0.05$) in variance.

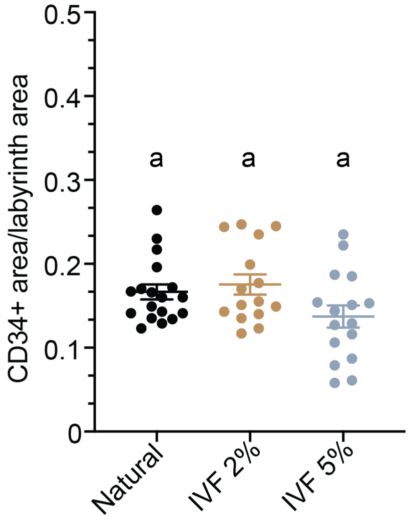


**Supplemental Figure 3: Additional gestational outcomes.**

Quantification of CD34+ positive staining in E12.5 placentas as a percentage of total labyrinth area using ePathology software. n ≥ 15 concepti from each group were used for CD34 quantification. Data are depicted as ± SEM and significant statistical differences between groups were determined via ordinary one-way ANOVA followed by Tukey's multiple comparisons test. Groups with different letters denote significant differences between groups (adjusted $P < 0.05$); same letters indicate no difference was detected.

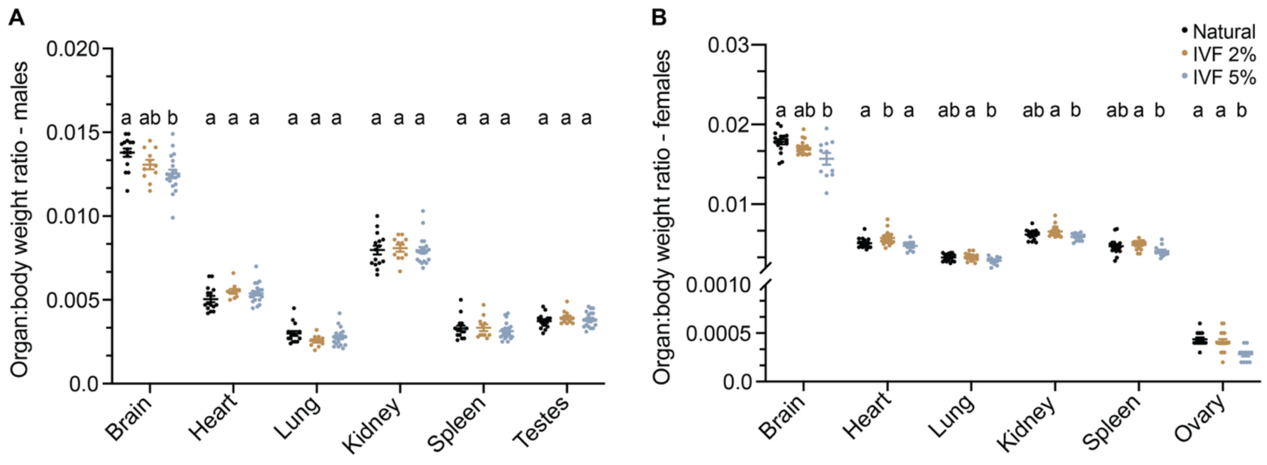


**Supplemental Figure 4: Organ to body weight ratios in 12-weeks-of-age offspring.** Organ to body weight ratios for **A**, male, and **B**, female offspring. n ≥ 11 male and female offspring were used in **A** and **B**. Data are depicted as ± SEM and significant statistical differences between groups were determined via ordinary one-way ANOVA followed by Tukey's multiple comparisons test. Groups with different letters denote significant differences between groups (adjusted $P < 0.05$); same letters indicate no difference was detected.
